# Regulation of Chromatin Architecture by Transcription Factor Binding

**DOI:** 10.1101/2023.09.26.559535

**Authors:** Stephanie Portillo-Ledesma, Suckwoo Chung, Jill Hoffman, Tamar Schlick

## Abstract

Transcription factors (TF) bind to chromatin and regulate the expression of genes. The pair Myc:Max binds to E-box regulatory DNA elements throughout the genome, controlling transcription of a large group of specific genes. We introduce an implicit modeling protocol for Myc:Max binding to mesoscale chromatin fibers to determine TF effect on chromatin architecture and shed light on its mechanism of gene regulation. We first bind Myc:Max to different chromatin locations and show how it can direct fiber folding and formation of microdomains, and how this depends on the linker DNA length. Second, by simulating increasing concentrations of Myc:Max binding to fibers that differ in the DNA linker length, linker histone density, and acetylation levels, we assess the interplay between Myc:Max and other chromatin internal parameters. Third, we study the mechanism of gene silencing by Myc:Max binding to the Eed gene loci. Overall, our results show how chromatin architecture can be regulated by TF binding. The position of TF binding dictates the formation of microdomains that appear visible only at the ensemble level. On the other hand, the presence of linker histone, acetylations, or different linker DNA lengths regulates the concentration-dependent effect of TF binding. Furthermore, we show how TF binding can repress gene expression by increasing fiber folding motifs that help compact and occlude the promoter region. Importantly, this effect can be reversed by increasing linker histone density. Overall, these results shed light on the epigenetic control of the genome dictated by TF binding.

## 1 INTRODUCTION

Eukaryotic genomes are compactly packaged inside the cell nucleus by wrapping ∼147 bp of DNA around an octamer of histone proteins. This first level of organization establishes the nucleosome, the chromatin basic repeating unit (1). Chains of nucleosomes form chromatin fibers that undergo additional folding that increases compaction. This folding can be aided by the binding of proteins. For example, CTCF and cohesin execute loop extrusion to form Topologically Associated Domains or TADs (2). Linker histones (LH) bind to the nucleosome at the entry/exit DNA sites to compact chromatin fibers and regulate their architecture (3, 4). Finally, transcription factors (TF) bind to regulatory DNA elements, modulating chromatin architecture and accessibility and, thereby gene expression (5).

Myc is a TF with a C-terminal basic helix-loop-helix leucine zipper (bHLHZip) motif that has DNA-binding activity and can establish protein-protein interactions (6). Myc regulates a large number of cellular processes, such as proliferation, growth, differentiation and pluripotency, metabolism, and apoptosis (7).

Myc heterodimerizes with another bHLHZip protein, Max, to form the Myc:Max complex that binds to E-box (5^′^-CACGTG-3^′^) regulatory DNA elements throughout the genome to control transcription (8). Myc:Max heterodimers can further form bivalent heterotetramers, allowing them to bring together sequence-distant regions of the genome (9). The heterotetramer assembles in a head-to-tail way through the individual leucine zippers of each heterodimer, resulting in the formation of an antiparallel four-helix bundle. This interaction between Myc and Max is essential for gene transcriptional repression (10). Several Myc-regulated genes contain multiple E-boxes within promoters that are usually separated by 100 bp (11). The binding of bivalent Myc:Max to such regions can create loops in the chromatin and allow the cooperative regulation at promoters and enhancers containing multiple E-boxes.

Efforts to model TF binding to chromatin have been made at various spatial levels. At the nucleosome level, all-atom and coarse-grained molecular dynamics simulations have been applied to study the binding of the pioneer TF Oct4, alone (12) or with Sox2 (13), to single nucleosomes. At the kb or chromosome level, TF binding has been introduced in polymer models as beads that can reversibly interact with other proteins or with the DNA (14, 15). Although the implicit binding of high-mobility group (HMG) proteins to chromatin fibers has been studied at nucleosome resolution (16), to the best of our knowledge, the effect of TF binding on chromatin architecture has not been studied at this resolution level.

Here, we perform mesoscale chromatin simulations at nucleosome resolution under the implicit binding of the Myc:Max complex to study this complex’s effect on microdomain formation (TAD-like structures at the kb level), its interplay with other chromatin elements (e.g., LH, histone tail acetylation, and linker DNA length), and its effect on gene folding. Overall, we find that depending on the linker DNA length, the TF binding position dictates the folding of fibers and the formation of microdomains. The extent of the effect of TF binding on chromatin compaction and overall shape depends on the presence of LH and histone acetylation, and on the length of linker DNA. While long linker DNAs and high acetylation levels allow higher TF concentrations to further compact the chromatin fibers, short linkers and LH impose structural restraints that limit the effect of TF binding. Finally, we show that the binding of Myc:Max can repress the expression of the gene Eed by increasing fiber folding and compaction, in a way that occludes the promoter region; interestingly, this repression can be reversed with high LH density.

Overall, our study provides new evidence on the regulation of chromatin architecture by TF binding and how such effects are regulated by chromatin composition and epigenetic marks.

## 2 METHODS

## 2.1 Chromatin Mesoscale Model

Briefly, our chromatin mesoscale model contains coarse grained elements that define the nucleosome cores, linker DNA, histone core tails, and LH (17–20) (Figure 1A). In particular, cores composed of 8 histone proteins and ∼147 bp of DNA are described as rigid cylinders with 300 Debye-Hückel charges distributed on their irregular surface calculated by the DiSCO algorithm (21). Linker DNA connecting nucleosomes is treated with a combined worm-like chain and bead model (22) based on the charged colloidal cylinder approach derived by Stigter (23) where each bead has a salt dependent charge and a resolution of ∼9 bp. Histone tails (18) and LHs (4, 24) are coarse grained as 5 residues per bead with the Levitt-Warshel united-atom protein model. The charges on histone tails beads are calculated as the sum of the five amino acids that compose the bead, whereas charges on LH beads are determined by the DiSCO algorithm. Acetylation of histone tails is modeled by alternatively folded rigid tails with altered force constants for stretching and bending with energies 100 times larger than wildtype tails (25). Tail coordinates are determined separately for wildtype and acetylated tails, by swapping coordinates when a change occurs. The new tail configuration after the swapping is subject to a standard Metropolis acceptance/rejection criterion based on the changes in local electrostatic energy (17).

**Figure 1:**
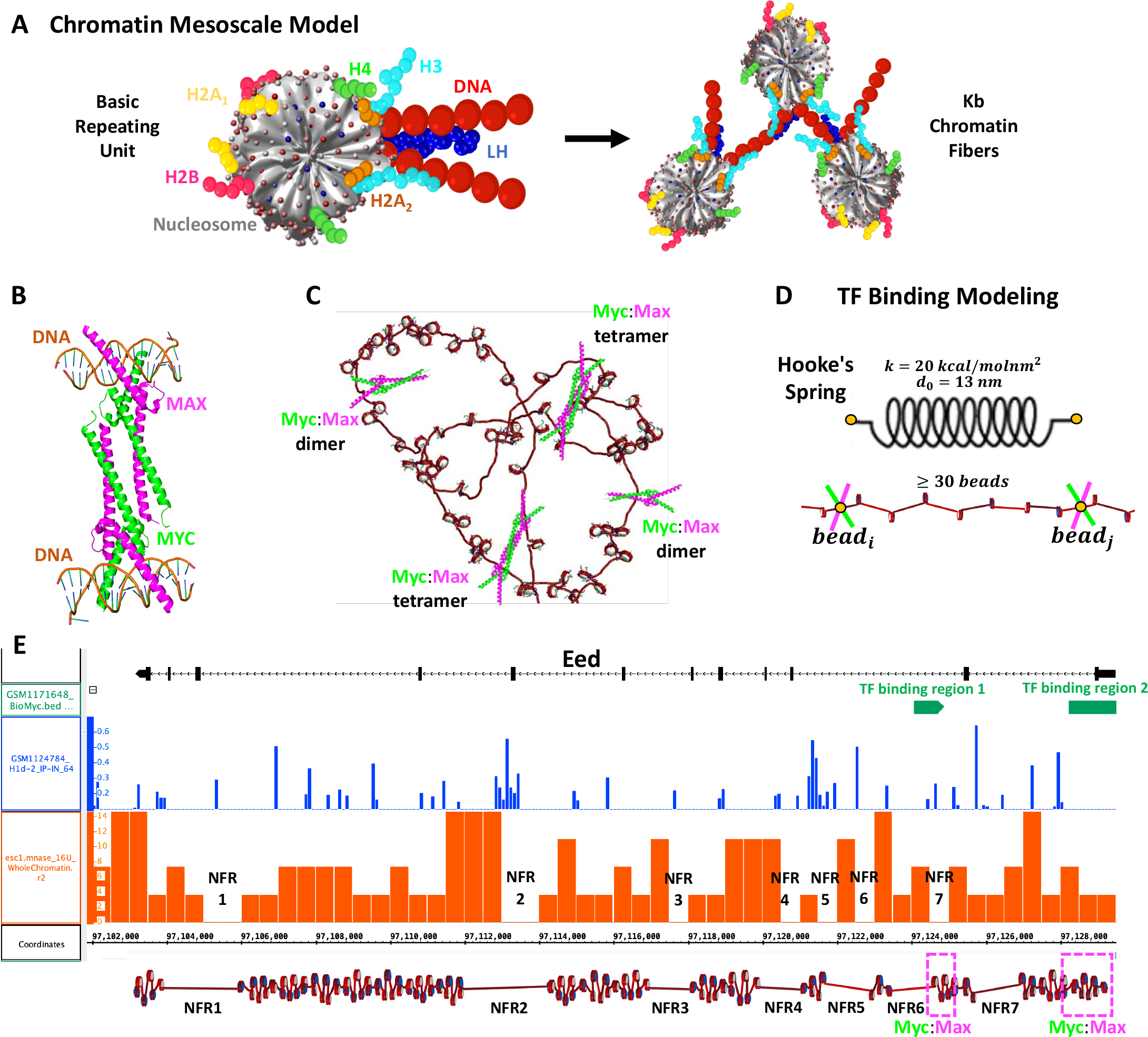
Our chromatin mesoscale model and modeling of Myc:Max binding to chromatin and the Eed gene. **A)** Coarse grained chromatin elements (nucleosome core, linker DNA, histone tails, and LH) form chromatin fibers at the kb level. **B)** Crystal structure of Myc:Max forming a heterotetramer that binds to sequence-distant DNA regions. **C)** Schematic representation of Myc:Max heterodimers and heterotetramers. **D)** Illustration of our implicit modeling of Myc:Max binding to chromatin fibers: distance constraints following Hooke’s spring law are enforced when two linker DNAs with Myc:Max bound are in close spatial proximity (20 nm) and at at least 30 linker DNAs apart. **E)** Design of the Eed gene loci using Mnase-seq data to position NFRs (orange track) and Chip-seq data to position LHs (blue track) and Myc:Max binding regions (green track).

The potential energy function of the model includes stretching, bending, and twisting terms for the linker DNA (*E*_*S*_, *E*_*B*_, *E*_*T*_), stretching and bending terms for histone tails (*E*_*tS*_, *E*_*t B*_) and LHs (*E*_*lhS*_, *E*_*lhB*_), and excluded volume (*E*_*V*_) and electrostatic (*E*_*C*_) terms for all beads as:

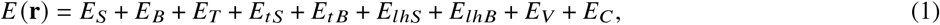

where **r** is the collective position vector.

Further details on the method parameters can be found in (17–20).

### 2.2 TF Binding Modeling

Based on the crystal structure (PDBID 1NKP) of the Myc:Max complex (Figure 1B) showing that heterotetramers can bind to sequence-distant regions, we simulate the binding of Myc:Max implicitly (Figure 1C) by adding restraints between two genome loci *b*_*i*_ and *b* _*j*_ (Figure 1D). In particular, a harmonic energy penalty of the form:

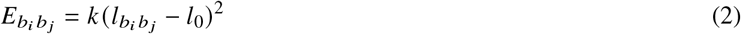

is applied to two target DNA linker beads. Here *l*_0_ is selected as 13 nm based on the distance between the two DNA chains in the crystal structure, and *k* is set to 20 *k cal*/*mol nm*^2^ as this value ensures that the energy penalty is not strong enough to produce overlapping of DNA beads or cores, but small compared to the total energy of the system.

During each trajectory, two DNA beads that have TF bound can engage in a constraint if the distance between them is less than 20 nm and if they are separated by at least 30 beads. If during the simulation, two beads that were engaged in a constraint separate more than 20 nm, the constraint is eliminated.

### 2.3 Systems

#### TF binding to different regions

To determine the effect of TF binding location on chromatin architecture, we simulate 50-nucleosome fibers without LH and histone acetylation, and with short, medium, or long linker DNA lengths, such as 26, 44, and 62 bp. Additionally, because real-life chromatin fibers are non unifrom, we also simulate a Life-Like fiber with linker DNAs that follow the distribution of linkers found in mouse embryonic stem cells (mESCs) (26): 30% for 26 bp, 17% for 35 bp, 15% for 44 bp, 13% for 53 bp, 9% for 62 bp, 7% for 70 bp 7, and 9% for 80 bp, as determined in our previous work (3).

For each system, we study four different binding patterns (see Figure 2A):

**Figure 2:**
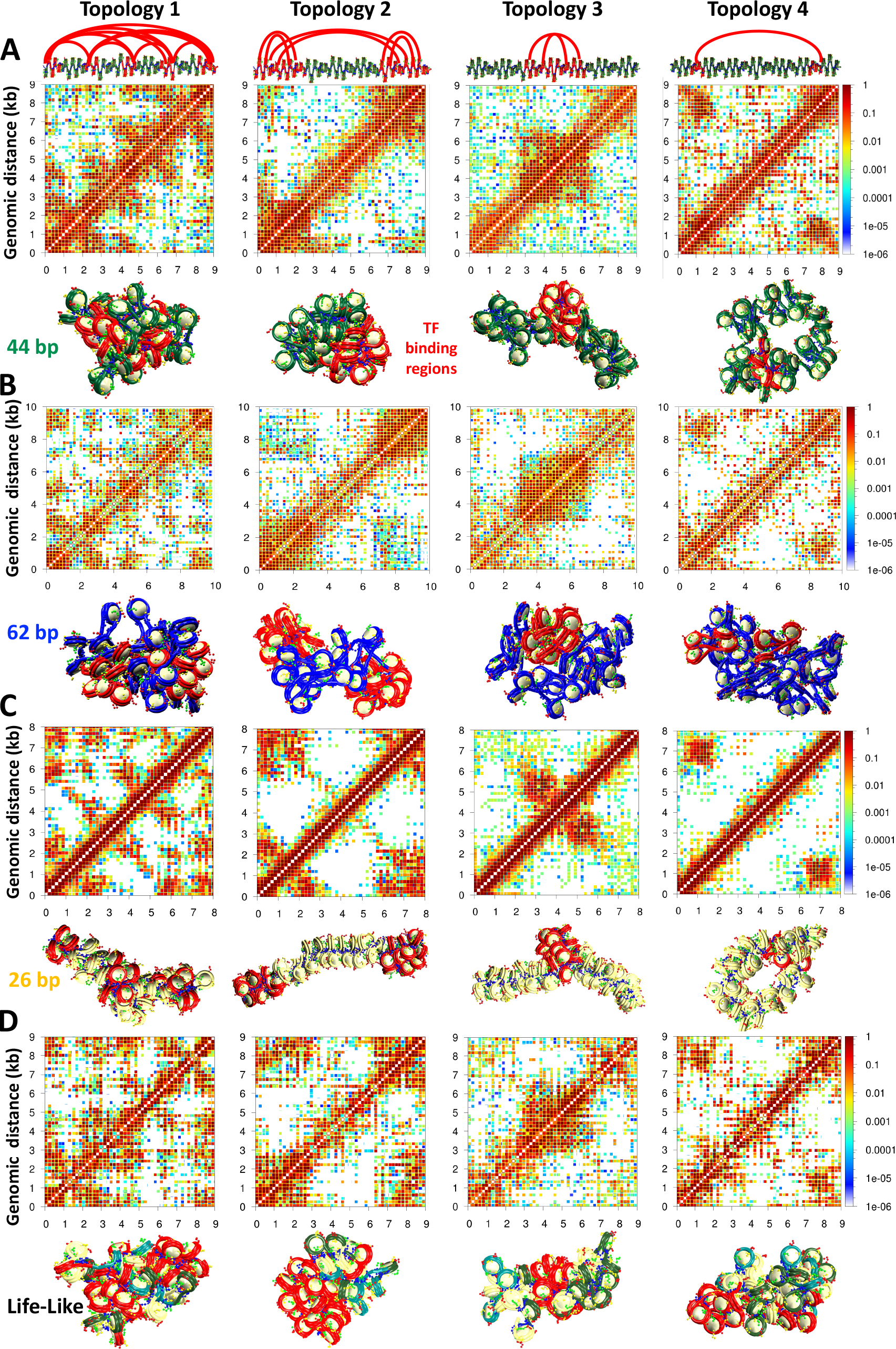
TF binding location drives formation of microdomains. 50-nucleosome chromatin fibers with: **A)** 44 bp linkers, **B)** 62 bp linkers, **C)** 26 bp linkers, and **D)** Non uniform linkers simulated with 4 different TF topologies. At the top, for the 44 bp system, we show an ideal zigzag fiber coloring in red the DNA with TF binding regions. Arcs show the possible binding geometries. The binding positions and geometries that define each topology are the same in all systems. For each system, we show the cumulative contact map calculated from 10 independent trajectories and a representative fiber structure also showing in red the TF binding regions. Additional representative structures are shown in Figures S2, S3, S4, and S5.

1. Topology 1: five equally distributed binding regions that occupy each the linker DNA of 5 consecutive nucleosomes
2. Topology 2: two binding regions, one at the beginning and one at the end of the fiber, that occupy each the linker DNA of 15 consecutive nucleosomes
3. Topology 3: one binding region located in the middle of the fiber that occupies the linker DNA of 15 consecutive nucleosomes
4. Topology 4: two binding regions, one located close to the beginning and another one located close to the end of the fiber, that occupy each the linker DNA of 5 consecutive nucleosomes

Note that by binding region we mean a region whose linker DNA beads are marked by one Myc:Max dimer.

#### TF binding at increasing concentrations

To determine the interplay between TF binding and chromatin internal parameters like LH, histone acetylation, and linker DNA length, we study TF binding at increasing concentrations such as 0, 25, 50, 75, and 100%. In particular, we study 50-nucleosome fibers of 26, 35, 44, 53, 62, 70, and 80 bp linker DNAs, as well as Life-Like fibers in three conditions:

1. Wildtype, systems have no LH and no acetylatation
2. +LH, systems have a density of 1 LH/nucleosome
3. +Acetylation, systems have four histone acetylation islands located at nucleosomes 6 to 11, 17 to 22, 28 to 33, and 39 to 44

In these three conditions, TFs are allowed to bind to any linker DNA region. TF concentration is calculated as the percentage of linker DNA beads that can bind TF.

#### Gene repression by Myc:Max binding

In the mm9 mouse genome assembly, Eed is located on chr7:97,103,164-97,129,486, occupying ∼26 kbp. To build Eed, Mnase-seq data (GSM2083107) (27) of mESCs were used to position nucleosome free regions (NFRs). In particular, the data were downloaded in bedgraph format from the Genome Omnibus Expression (GOE) repository and loaded into the UCSC Genome Browser without further processing. NFRs were visually inspected and identified as genomic regions with absence of signal. Details of MNase-seq data processing can be found at the GEO site where the data are deposited. As shown in Figure 1E, 7 NFRs can be identified. NFRs 1 and 2 are long and we model them with ∼ 350 bp, whereas NFRs 3 to 7 are short, and we model them with ∼ 200 bp. Similar to the Life-Like fibers, the linker length distribution of mESCs (26) was used to determine the distribution of non uniform linker lengths, and the nucleosome repeat length (NRL) typical of mESCs, 189 bp (28), was used to calculate the number of nucleosomes in the 24 kbp fiber (length of the fiber without the NFRs), obtaining 129 nucleosomes. In the Eed gene, two binding regions for Myc:Max have been detected with Chip-seq experiments (GSE48175) (29) (Figure 1E). One between 97,128,221 and 97,129,793 bp, and a second one between 97,124,070 and 97,124,896 bp. Thus, in the Eed model, these regions are defined as TF binding regions 1 and 2 (Figure 1E). Similarly, LH positions were determined based on Chip-seq data (GSE46134) (30) where peaks with frequencies at least 10% of highest frequency peak were selected (Figure 1E), producing a density of 0.37 LH/nucleosome, close to the average LH density of 0.5 found in mESCs (28). Histone acetylation was not introduced in the model, as analyzed Chip-seq data (31–33) show no acetylation in the Eed region. To determine the role of LH in the Eed activation mechanism, we additionally simulate the Eed system (with and without TF) with LH density *ρ* = 0.8 (LHs randomly distributed), as found in mouse somatic cells (28). The list of linker DNAs, LH positions, and Myc:Max positions are detailed in the Supporting Information, Table S1.

### 2.4 Simulation and Analysis

Each system is sampled with Monte Carlo (MC) simulations. For the 50-nucleosome systems, we use 50 million MC steps and 10 independent replicas. For the Eed system of 129 nucleosomes, we use 70 million MC steps and 20 independent replicas. All replicas are started from different random seeds and a DNA twisting angle of 0, +12, or -12 degrees to mimic natural variations (34). Frames are saved every 100,000 MC steps.

The MC moves include a global pivot move that selects the shorter fiber end passing through a random axis and rotates it, local translation and rotation moves of DNA beads and cores, and local translation of LH beads (17). Histone tails are sampled with a bead-by-bead regrowth move by the Rosenbluth scheme (18).

Trajectory convergence is monitored by the system energy and global (end-to-end distance and sedimentation coefficient) and local (nucleosome triplet angle) parameters (Figure S1) over the entire ensemble (10 trajectories for each 50-nucleosome system and 20 trajectories for the Eed gene). For the 50-nucleosome systems, the last 10 million steps, or 100 structures, of each of 10 trajectories are used to create ensembles of 1000 structures per system. For the Eed gene systems, we create ensembles of 2000 structures with the last 10 million steps of 20 trajectories. These ensembles are analyzed to calculate fiber packing ratio, sedimentation coefficients, radius of gyration, and volume and area of the promoter region.

The fiber packing ratio is calculated as:

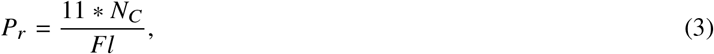

where *N*_*C*_ is the total number of nucleosomes and *Fl* is the fiber length calculated by defining the fiber axis with a cubic smoothing spline interpolation to the nucleosomes *x, y*, and *z* coordinates; see details in the supporting information of (3).

Sedimentation coefficients are calculated as:

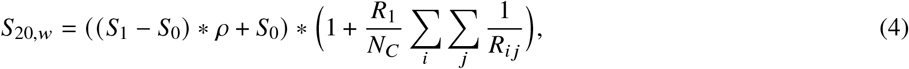

where *S*_0_ and *S*_1_ are the sedimentation coefficients of a mononucleosome without LH (*S*_0_ = 11.1 *S*) (35) and with LH (*S*_1_ = 12 *S*) (36), respectively, *ρ* is the LH density on the fiber, *R*_1_ is the radius of a nucleosome (*R*_1_ = 5.5 nm), *N*_*C*_ is the number of nucleosomes in the chromatin fiber, and *R*_*i j*_ is the distance between the nucleosomes *i* and *j*.

The radius of gyration, which describes the overall dimension of the chromatin fiber, is measured as the root mean squared distance of each nucleosome from the center of mass according to:

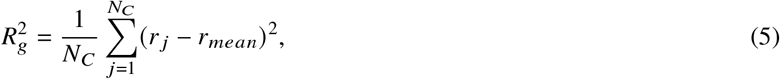

where *N*_*C*_ is the number of nucleosomes, *r* _*j*_ is the center position of the nucleosome core j, and *r*_*mean*_ is the average of all core positions

The volume and area of the Eed promoter region that overlaps with the TF binding region 1 are calculated using the alphashape function of Matlab. With alphaShape, a bounding area or volume is created to envelop the nucleosome coordinates. We use an alpha value of 100 to create a loose shape. In particular, we use the x,y (for area) or x,y,z (for volume) coordinates of nucleosomes 123 to 129 as vertices, as they are located in the promoter region.

Interactions among nucleosomes are calculated every 100,000 MC steps for each trajectory, normalized across all trajectory frames, and summed to create a contact map that we plot at logarithmic scale. Two nucleosomes *i* and *j* are considered to be in contact if any element of nucleosome *i*, such as core, tails, or linker DNA, is less than 2 nm from any element of nucleosome *j*. These matrices are calculated in both bp and nucleosome resolution.

Internucleosome interaction matrices at nucleosome resolution are decomposed into one-dimensional plots that depict the magnitude of *i, i* ± *k* interactions, or contact patterns, as follows:

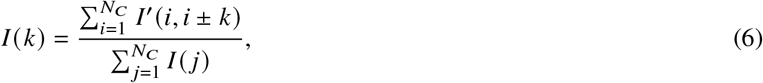

where *N*_*C*_ is the number of nucleosomes, *I* is the internucleosome interaction matrix, and *k* is the number of nucleosomes between cores *i* and *j*.

To estimate the number of microdomains in each system, we convert the interaction matrices in nucleosome resolution to distance matrices by calculating the inverse of each element in the matrix (*d* = 1/ *f r equency*) and perform a clustering analysis with the DBSCAN algorithm (37). To define the clusters, we set the parameter *minpoints* as 5 for every system, and the radius of search, *ϵ*, as 3 for the 62 bp system, 2 for the 44 bp and life-like systems, and 1.4 for the 26 bp system.

## 3 RESULTS

### 3.1 TF Binding Mediates the Formation of Microdomains

Protein binding to chromatin fibers has shown to produce regions of enriched contacts in Micro C or Hi-C maps. For example, loops associated with chromosome domain boundaries overlap with CTCF binding regions (38). Similarly, enrichment of Hi-C contacts can be used as a reporter of the strength of interaction between a pair of TF binding sites (39). Finally, binding of TFs, cofactors, or chromatin modifiers produces fine-scale domains, smaller than TADs, that can be identified as stripes and dots in Micro C maps (40).

To better understand how TF binding position affects chromatin architecture, we simulate 50-nucleosome uniform fibers of 26, 44, and 62 bp linker DNAs, as well as 50-nucleosome Life-Like fibers with TF binding in four different scenarios (Figure 2). Interestingly, contact maps of fibers with both medium (44 bp) (Figure 2A) and long (62 bp) (Figure 2B) linker DNAs show clear regions of high-frequency contact, or microdomains, as a result of the TF binding, that are dependent on the TF binding positions. In fibers with short linkers (Figure 2C), like 26 bp, the formation of microdomains is less clearly identified in the contact maps. Namely, high-intensity regions close to the diagonal are diffuse and not as well defined as in the fibers with 44 and 62 bp linkers. See for example topologies 2 and 3 in panels A, B, and C of Figure 2. On the other hand, other topologies, like topology 1 and 4, allow identification of microdomains. The 44 and 62 bp linker fibers are less compact and more globular than the 26 bp linker fiber (20). Additionally, these fibers are more sensitive to changes in salt concentration and presence of LH (3, 20). The presence of more diffuse microdomains in 26 bp fibers can be explained by the rigidity of short-linker fibers in which the linker DNA dictates fiber architecture and produces a highly bent 10-nm ladder-like form (19) with low sensitivity to external and internal parameters, such as LH binding (3, 20, 41). Thus, longer linker DNAs that give chromatin fibers more flexibility, facilitate their folding modulation by TF binding.

Similar to the 44 and 62 bp uniform systems, microdomains emerge in ensemble-based contact maps of Life-Like fibers (Figure 2D). Due to the fiber polymorphism triggered by variations in the linker DNA (19), the microdomains are slightly more diffuse than in the 44 and 62 bp uniform fibers (Figure 2A and B). Thus, TF binding positions define the organization of Life-Like chromatin fibers, but the irregularity of the fibers smooths out this effect compared to uniform systems; fibers that combine short and long linkers form fluid heterogeneous bent ladders that accommodate TF binding in a more relaxed way than compact 30-nm regular forms.

As shown in Figure S6, these microdomains can also be identified by a clustering analysis of the contact map in nucleosome resolution. Fibers with 44 and 62 bp linkers show clear TF-topology-dependent nucleosome clusters. In contrast, for the 26 bp and Life-Like fibers, the TF dependency of the nucleosome clusters is not as clear. These results support the notion that microdomains are more diffuse in these two systems due to fiber rigidity or polymorphism, respectively. Similarly, internucleosome contact plots (Figure S6) show some TF binding position variations in medium and long-range interactions for the 44 and 62 bp fibers, but less variations for the 26 bp and Life-Like systems. See for example the plots corresponding to Topology 1 of each system where the 44 and 62 bp fibers reveal 4 clear high-frequency regions at medium and long-range interactions that are not as clear in the 26 bp and Life-Like systems.

Importantly, the contact maps that reveal microdomains are ensemble-based maps in which the contacts from 10 independent trajectories are summed. These maps are analogous to experimental Hi-C maps obtained from a population of cells. On the other hand, the contact maps corresponding to single trajectories, equivalent to single-cell Hi-C maps, do not show clear formation of microdomains. As we see from Figure 3 for the 62 bp system with 5 TF binding regions (Topology 1), checker-board patterns in the ensemble-based map are a product of TF binding not discernible in individual contact maps obtained from single trajectories. These results agree with experimental data showing that cohesin dictates the preferential positioning of boundaries at CTCF sites to produce ensemble TAD structures (42). Here, such microdomain structures emerge naturally from our tagging of TF binding regions.

**Figure 3:**
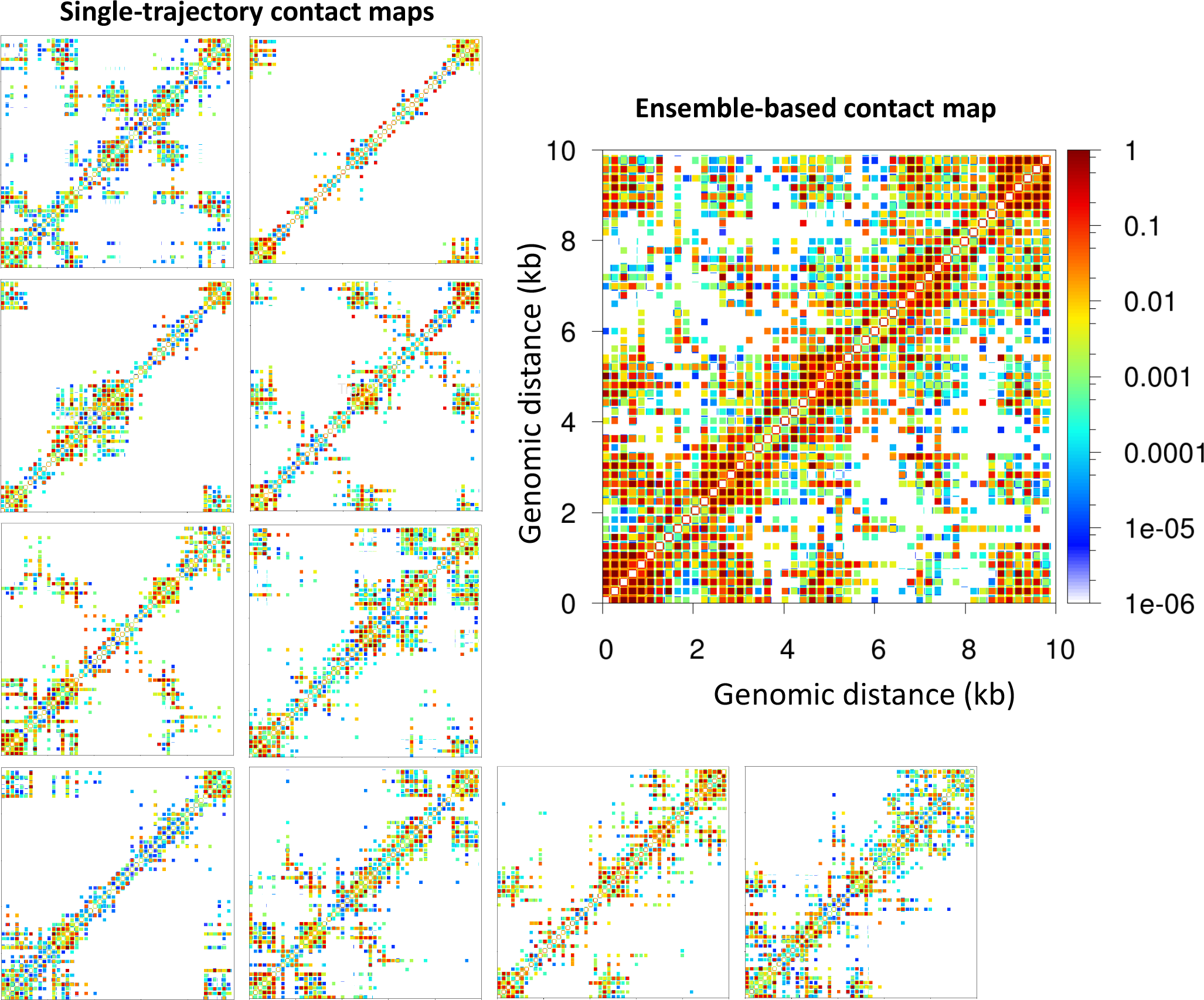
Microdomains emerge only from ensemble-based contact maps. The 10 single-trajectory contact maps for the 62 bp system Topology 1 (5 TF binding regions) at left reveal various microdomains. The large ensemble-based contact map at right obtained by summing the 10 individual contact maps reveals all possible microdomains contacts.

### 3.2 TF binding Effect is Regulated by Linker DNA length, Histone Acetylation, and LH density

As we have shown previously, histone acetylation affects chromatin compaction (25), size and compaction of nucleosome clutches (43), and can produce the segregation of domains (44). Similarly, LH density controls chromatin higher order folding and compaction (3, 4, 45), and the size and compaction of nucleosome clutches (43). In addition, LH binding and tail acetylation act cooperatively to direct fiber folding (17).

To investigate how TF binding and other chromatin regulators, such as LH and tail acetylation act together to modulate chromatin architecture, we study TF binding in presence of LH and tail acetylation for fibers with different linker DNAs. In particular, we determine TF saturation curves for fibers with uniform linkers (26, 35, 44, 53, 62, 70, and 80 bp), and for non uniform Life-Like fibers by measuring both packing ratios and sedimentation coefficients at increasing TF concentration (percentage of linker DNA beads that can bind TF), from 0 to 100%.

As we see from Figure 4A and Figure S7, when the linker DNA is short, such as 26 and 35 bp, TF binding does not increase the packing ratio of the fiber. However, the sedimentation coefficient increases significantly from 0 to 25% [TF], and then remains steady. This indicates that TF binding saturation occurs at a concentration of 25% and that higher TF concentrations do not affect fiber folding. Namely, fibers cannot fold onto a more compact form due to excluded volume interactions. As clearly shown by the fiber configurations of the 26 bp system (Figure 5), while in absence of TF the fiber has a ladder-like extended structure (short linkers), it folds over itself when TF is bound, explaining the higher sedimentation coefficients. On the other hand, for medium and long linker fibers, both packing ratios and sedimentation coefficients increase upon TF binding (Figure 4A). The longer linkers allow nucleosomes to approach closer upon TF binding, increasing the packing ratio. Additionally, the longer the linker DNA, the larger the change in packing ratio upon TF binding at a concentration of 25%. Similar to the short linker fibers, TF binding also produces a more globular fiber (Figure 5) with higher sedimentation coefficient (Figure 4A).

**Figure 4:**
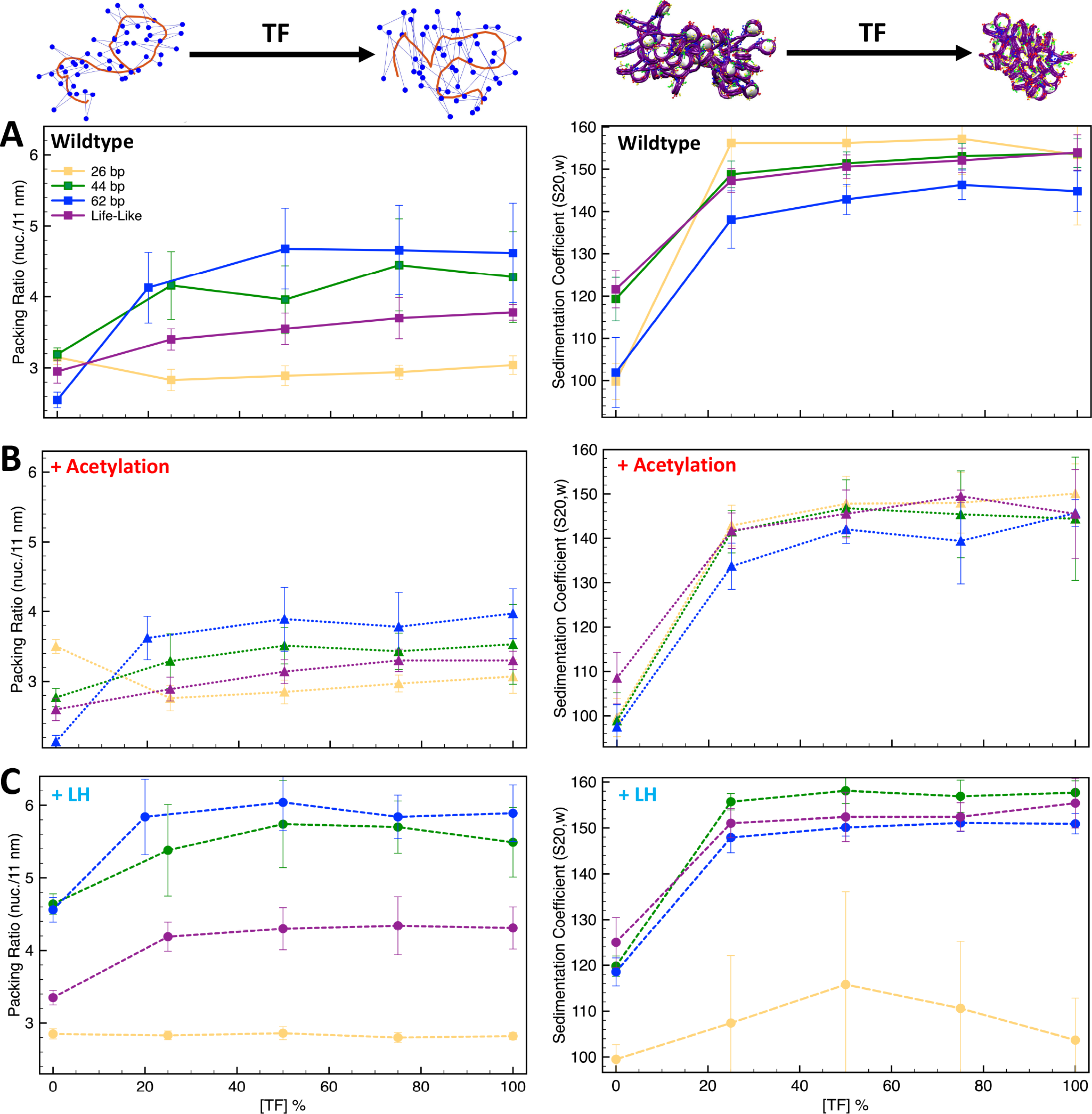
TF saturation curves are affected by histone acetylation and LH. The graphs **A–C** show packing ratios and sedimentation coefficients as a function of TF concentration for the 26 bp, 44 bp, 62 bp, and Life-Like fibers in different conditions: **A)** systems without LH and acetylation; **B)** systems with two acetylation islands; and **C)** systems with LH density *ρ* = 1. At top left, we show the fiber axis (red trace) and position of nucleosomes (blue dots) for a 70 bp chromatin fiber to illustrate the increase of packing ratio (number of nucleosomes per 11 nm of fiber length) upon TF binding. At top right, we show a 70 bp linker chromatin fiber to illustrate the decrease of chromatin global size upon TF binding. Results including uniform systems with 35, 53, 79, and 80 bp are shown in Figure S7.

**Figure 5:**
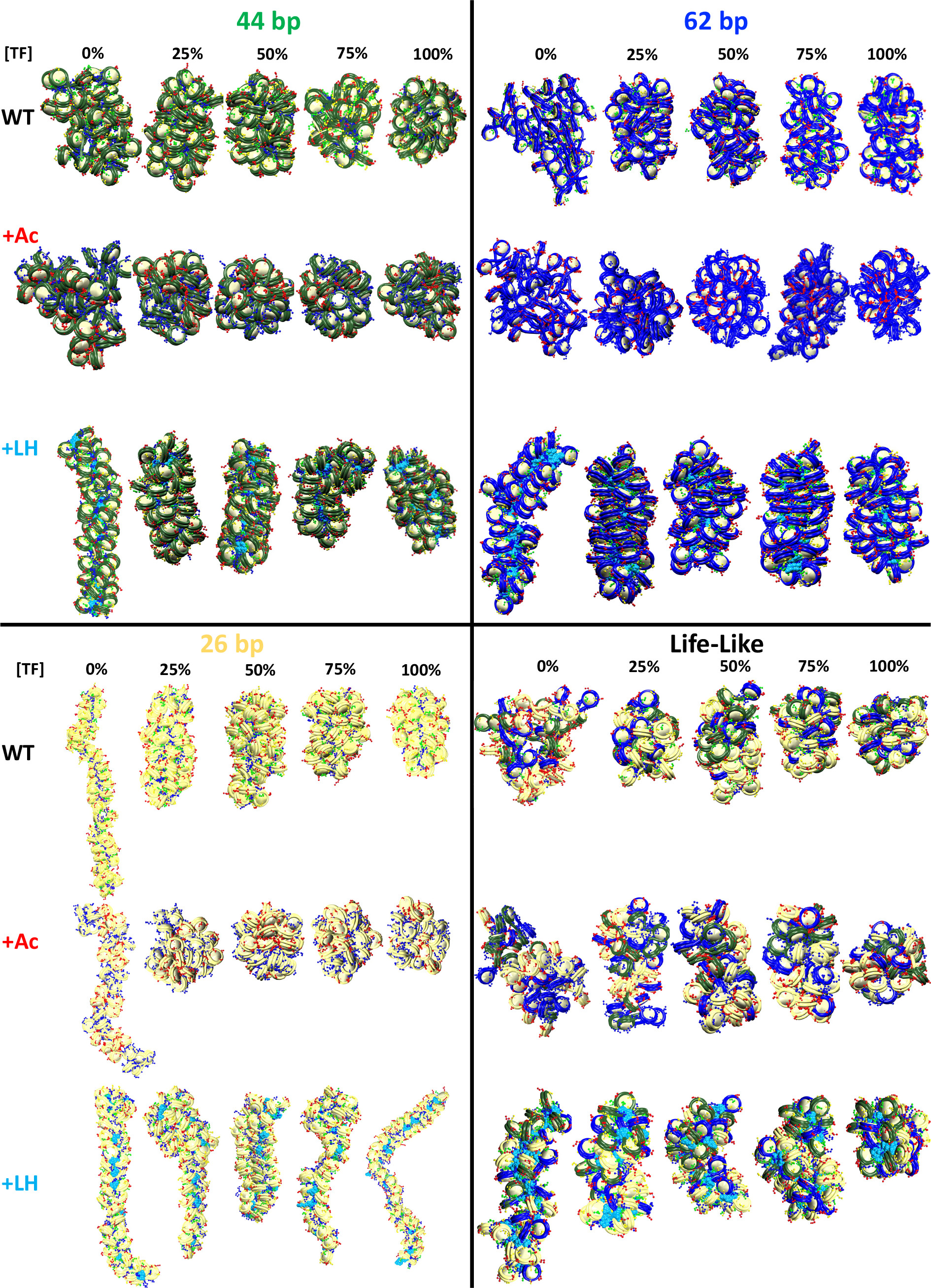
TF binding affects chromatin architecture. Chromatin uniform fibers of 44, 62, and 26 bp linkers, as well as non uniform Life-Like fibers at increasing TF concentration. WT shows structures of the wildtype systems (no acetylation and no LH). +Ac shows structures of the acetylated systems with acetylated tails drawn in red and wildtype tails in blue. +LH shows structures of systems with an LH density of 1 LH/nucleosome, with LHs drawn in cyan.

For Life-Like fibers, the effect on packing ratio and sedimentation coefficient is similar to that observed in the medium and long linker uniform fibers; fibers become more compact and globular upon TF binding (Figure 4A and Figure 5). However, in agreement with the trends observed for the TF binding regions, fluid Life-Like fibers appear to be less sensitive to the increase of TF concentration than the 44 and 62 bp systems.

When acetylation islands are incorporated into the chromatin systems (Figure 4B), the trends in packing ratios and sedimentation coefficients are similar to wildtype systems, but the extent of the TF binding effect is diminished. There is a smaller change in packing ratio when we increase TF binding from 0 to 25%. This is because histone acetylation disrupts internucleosome contacts and opens the chromatin fiber structure (25), which is opposite to the repressive effect observed by TF binding. In agreement, compared to wildtype systems, fiber structures (Figure 5) show a smaller change in global shape and compaction upon TF binding.

Similarly, when LH is bound to medium and long linker DNA fibers (Figure 4C), the effect of TF on chromatin compaction is diminished compared to the wildtype systems. In this case, LH is a chromatin compactor, like TF binding. Thus, LH and TF compete for compacting the chromatin fiber. At 0% TF, the packing ratio of the systems with LH are much higher than the wildtype systems due to LH binding. Thus, upon TF binding, the fiber packing ratio cannot increase as when no LHs are bound. Moreover, LH binding produces straighter and ordered structures (3), whereas TF binding produces more globular fibers. Thus, the effects of TF and LH on fiber global shape are in opposition. Upon TF binding, fibers with LH are less globular and more straight than the equivalent wildtype fibers (Figure 5).

For short linker fibers like 26 and 35 bp, as we saw in the wildtype system, no increase can be observed in packing ratio upon TF binding. However, very different trends are observed for the sedimentation coefficients of the 26 bp system. In contrast to the wildtype system (Figure 4A), TF binding does not increase the sedimentation coefficient (Figure 4C), and larger standard deviations are obtained, indicating more variability in the fiber structures that create the 1000-structure ensemble. This is because with LH, some TF dimer/dimer interactions cannot form in some of the trajectories. For example, for the 50% [TF] system, from the 10 independent trajectories, 4 reach the 50% [TF], 4 have 0% [TF], and 2 have less than 20% [TF], producing an effective [TF] in the ensemble of ∼22%. Thus, LH bound to short linker DNAs produces a fiber rigidity that impairs TF function. A similar effect has been seen for condensin and topoisomerase II binding (46), where LH depletion causes excess loading of these proteins to chromosomes, producing aberrant chromosomes.

Finally, for Life-Like fibers (Figure 4C), the trends in packing ratio and sedimentation appear similar to those of the wildtype systems (Figure 4A). Interestingly, the increase in packing ratio upon TF binding at a 25% concentration is slightly higher than for the wildtype system (0.85 vs. 0.45 increase). This indicates that simultaneous binding of LH and TF to Life-Like fibers is better accommodated than in uniform fibers. Thus, fiber fluidity triggered by variations of linker DNA (19) modulates the effect of protein binding on chromatin architecture.

### 3.3 TF Binding-Mediated Compaction as Possible Mechanism of Gene Locus Repression

Myc helps maintain cell pluripotency and inhibits differentiation by activating genes needed for pluripotency and repressing genes that trigger differentiation (29, 47). For example, the embryonic ectoderm development gene, Eed, expresses a protein member of the Polycomb-group (PcG) family required for silencing pluripotency genes upon embryonic stem cell (ESC) differentiation (48). Thus, in ESCs, Eed is repressed so that cell differentiation is suppressed. To determine the mechanism of gene repression in Eed by Myc:Max, we fold *de novo* the Eed gene, in the presence and absence of TF binding. To study Eed gene activation upon cell differentiation, we additionally simulate the folding of Eed with and without TF binding but increasing LH density (*ρ*) to 0.8 LH/nucleosome, the level observed in mouse somatic cells (28).

Figure 6A shows the ensemble-based contact map for the Eed gene with and without TF binding. We see that TF binding increases the number and frequency of internucleosome contacts. The density of the matrix increases by ∼9% upon TF binding. On the right upper corner of the contact maps, there is a clear difference: microdomains emerge as product of the TF binding. Contact maps obtained from single trajectories (Figure 6B) clearly show that TF binding increases hairpin motifs (medium-range contacts) and hierarchical loops (long-range contacts); these motifs are evident from diagonal and perpendicular regions to the main diagonal of the map (45). Such an increase in local loop interactions is also observed in the plot of contact frequency versus genomic distance (Figure S8) showing that while short local interactions slightly decrease upon TF binding, local loop interactions increase markedly. These folding motifs increase fiber compaction (Figure 6C). Fiber configurations show that upon TF binding, the TF binding regions are in close proximity, producing a more compact fiber.

**Figure 6:**
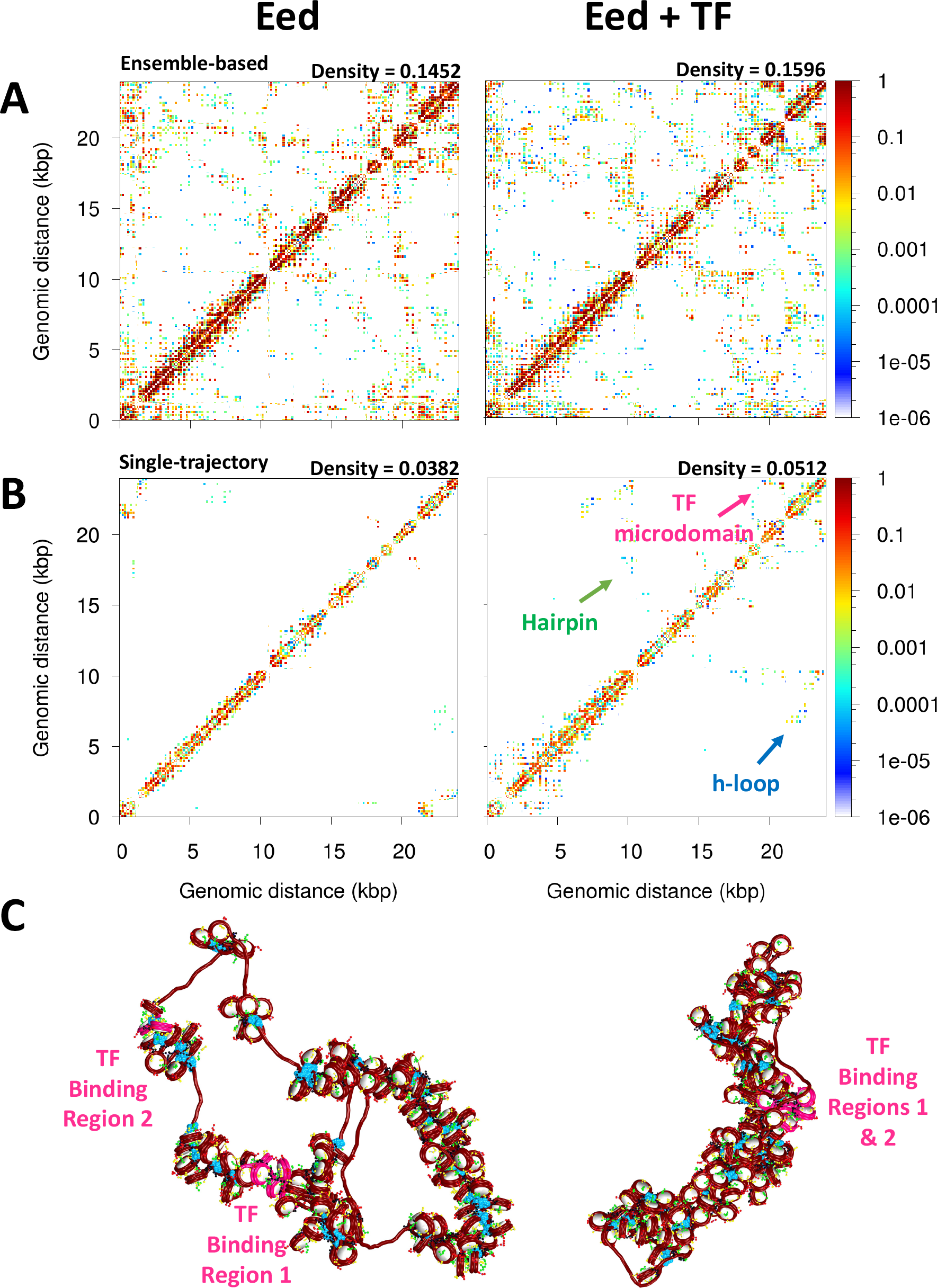
TF binding repress the Eed gene loci. **A)** Ensemble based nucleosome contact maps obtained from 20 independent trajectories of the Eed gene in absence and presence of TF binding. **B)** Nucleosome contact maps obtained from a single trajectory of the Eed gene in absence and presence of TF binding. **C)** Representative chromatin fibers of the Eed gene in absence and presence of TF binding. In magenta are shown the TF binding regions. LHs are shown in cyan.

Furthermore, fiber sedimentation coefficient increases, while radius of gyration decreases, upon TF binding (Table 1). Such radii of gyration measurements agree with experimentally and theoretically determined radii of gyrations for similar size fibers (49, 50) (e.g., our 62 ± 14 nm Rg is similar to the 43 ± 9 nm Rg predicted for 26 kb regions in Drosophila cells (50)). Significantly, we find that the TF binding region 1, which overlaps with the promoter region, becomes occluded, thus less accessible. As reported in Table 1, the area and volume of the promoter region are reduced ∼1.5 times upon TF binding. Thus, Eed mechanism of repression by Myc:Max involves the compaction and occlusion of the promoter region due to increased fiber folding. A less accessible promoter will impair gene transcription.

**Table 1:** Compaction parameters: Sedimentation coefficient and radius of gyration for the entire Eed system, and area and volume for the promoter region of Eed.

| | Fiber Rg (nm) | Fiber Sc ($S_{20,w}$) | Promoter Area (nm<sup>2</sup>) | Promoter Volume (nm<sup>3</sup>) |
| --- | --- | --- | --- | --- |
| Eed | $62 \pm 14$ | $162 \pm 19$ | $252 \pm 45$ | $1753 \pm 239$ |
| Eed + TF | $60 \pm 13$ | $171 \pm 18$ | $171 \pm 42$ | $1262 \pm 202$ |
| Eed + 0.8 LH | $61 \pm 9$ | $173 \pm 15$ | $196 \pm 47$ | $1447 \pm 123$ |
| Eed + 0.8 LH + TF | $57 \pm 8$ | $175 \pm 16$ | $210 \pm 46$ | $1420 \pm 104$ |

Upon cell differentiation, LH density increases to ∼0.8 LH/nucleosome (28), and Eed needs to be activated so the polycomb protein is expressed and the pluripotency genes repressed. Thus, to determine how this LH density increase upon cell differentiation might affect Eed repression and help its activation, we study the Eed gene loci with an LH density *ρ* = 0.8. When *ρ* increases, the fiber becomes more compact and less flexible (Figure 7 and Table 1) due to the repressive effect of LH binding (3, 24). This change in fiber compaction and flexibility impacts TF influence. Similar to the effect observed for the 26 bp system in Figure 4, the Myc:Max dimers cannot interact when the fiber becomes more rigid and straight due to LH binding. Indeed, from the 20 trajectories ran for this system, only one trajectory has dimers engaged, and thus, the systems with *ρ* = 0.8 with and without TF exhibit similar properties (Table 1), structures (Figure 7), and genomic contacts (Figure S9). These results implicate LH in the activation of genes by impairing the effect of repressing proteins.

**Figure 7:**
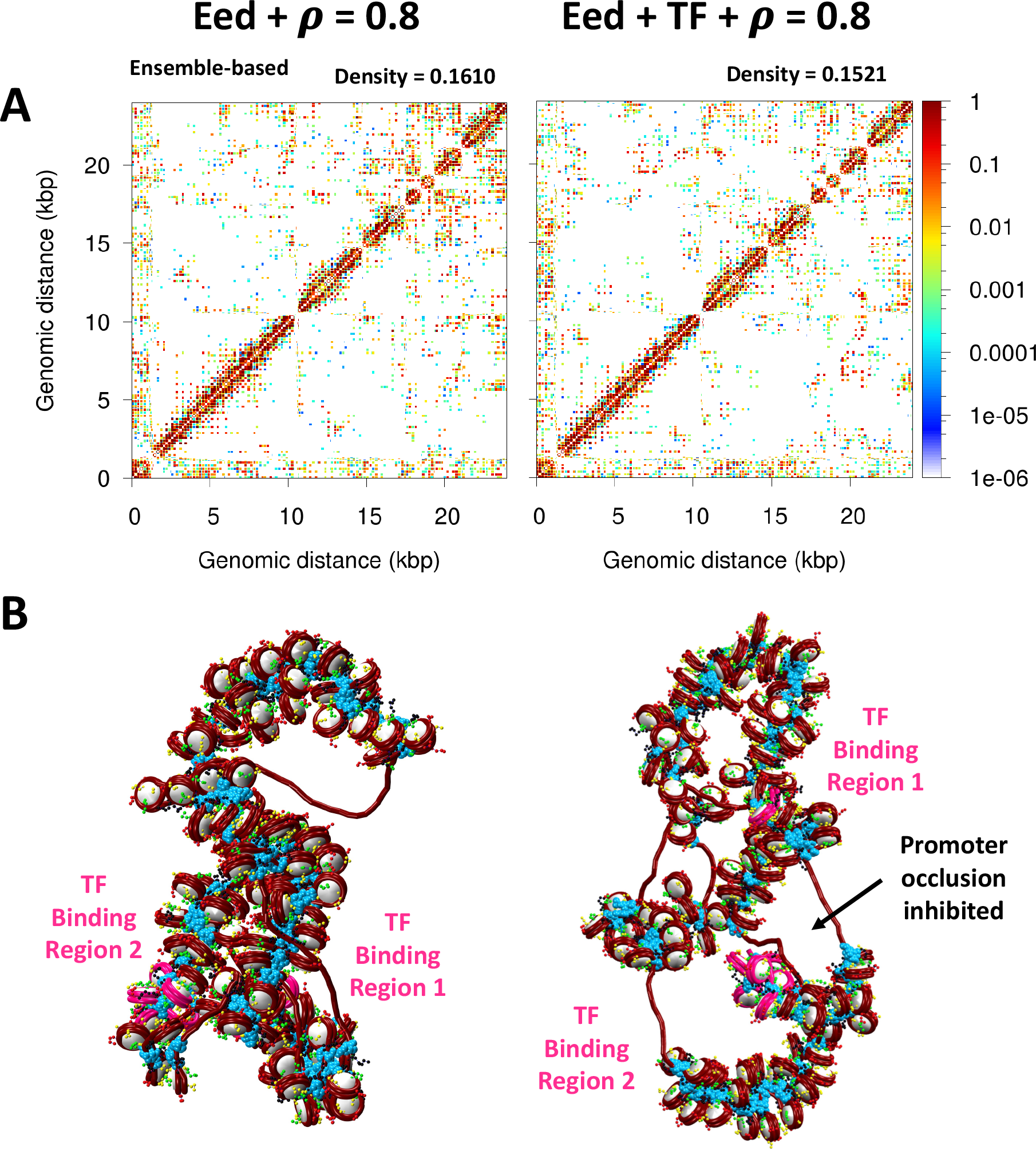
Activation of the Eed gene loci depends on LH density. **A)** Ensemble based nucleosome contact maps obtained from 20 independent trajectories of the Eed gene with an LH density of 0.8 LH/nucleosome in absence and presence of TF binding. **B)** Representative chromatin fibers of the Eed gene in absence and presence of TF binding showing that the two TF binding regions remain apart upon TF binding. In magenta are shown the TF binding regions. LHs are shown in cyan.

## 4 DISCUSSION

Our Monte Carlo simulations of chromatin fibers at nucleosome resolution with implicitly bound TFs demonstrate how these proteins can shape chromatin architecture and thus regulate gene expression. Our results on different TF binding regions (Figure 2) clearly show that TF binding creates microdomains that are visible in the ensemble-based contact maps; these microdomains are more clearly evident in, and enhanced by longer-linker DNAs. These results agree with experimental Hi-C maps (38–40) and polymer modeling studies (14, 51, 52) showing that the clustering of proteins that bridge genomic segments results in TAD-like structures. Our results showing that in fibers with short linkers microdomains are more diffuse due to geometrical restrictions agree with observations that silent chromatin has longer linker DNA than active chromatin (53). Longer linkers provide the chromatin fiber with the flexibility needed to fold and create loops that compact the structure and repress transcription. We showed this for metaphase chromatin where longer linkers and the absence of LH favor the formation of hierarchical loops that compact genes and chromosomes (45). Importantly, Life-Like fibers, representative of real-life chromatin, are sensitive to TF binding positions and form microdomains, although these are less defined than in uniform fibers due to the high plasticity and fluidity of the heterogeneous chromatin (19).

Similar to the effect on microdomain formation, we also suggest that, upon TF binding, fibers with short linkers (26 and 35 bp) do not change their packing ratio, whereas fibers with medium and long (44 to 80 bp) linkers, as well as Life-Like fibers, become more compact (Figure 4A). Thus, the role of linker DNA appears to be coupled to the effect that repressive proteins have on the chromatin structure. This agrees with recent cryo-EM results showing that long linker DNAs produce relaxed DNA trajectories that enable the binding of LH and produce transcriptionally silent heterochromatin regions, whereas short linker DNAs preclude the binding of LH and produce transcriptionally active chromatin (54).

Histone tails extend from the nucleosome core and interact with other nucleosomes and with the linker DNA (18). When tails are acetylated, they become globular, impairing internucleosome interactions and opening up the chromatin fiber (25). When we introduced histone acetylation, we saw that, for fibers with medium, long, and Life-Like linker DNAs, the effect of TF binding on chromatin compaction is reduced (Figure 4B). This is because the effects of acetylation and TF binding (Myc:Max in particular) on chromatin compaction and architecture are opposite. In line with these results, it has been reported that histone acetylation can prevent the binding of some pioneer TFs (55). Our results thus highlight how histone acetylation can counter and modulate the repressive effect of proteins that bind to chromatin.

When studying the effect of LH binding, we found that LH can impair the effect of TF binding on chromatin architecture (Figure 4C). Particularly affected were fibers with short linkers. The C-terminal domain of LH interacts with the linker DNAs to compact chromatin fibers (24). Thus, LH competes for the linker DNA with TF binding. Indeed, recent experiments have shown that although LH remains bound to the nucleosome dyad upon TF binding, the C-terminal domain unbinds from the linker DNA (56). These results highlight the influence of the LH C-terminal domain on regulating TF binding. LH binding produces a rigid and straight fiber (3) that is not flexible enough to be maneuvered by TF binding. Thus, although TF and LH binding might not be mutually exclusive as they do not share the same genomic binding sites, LH can indirectly impair TF function by affecting the chromatin fiber architecture. Interestingly, this effect seems to be less important in Life-Like fibers; these fibers better accommodate simultaneous LH and TF binding. Thus, fiber fluidity appears essential to allow the regulation of chromatin fibers by multiple protein binding.

Our folding of the Eed gene loci upon repression and activation revealed that TF binding produces the formation of microdomains that can occlude the promoter of Eed, repressing its transcription (Figure 6). However, this effect is eliminated by increasing the LH density (Figure 7), as the fiber flexibility does not allow for the DNA regions with TF bound to come together. Thus, TF and LH binding work together to regulate gene expression. This cooperation further underscores the idea that LH does not always act as a repressor of global transcription, and agree with results found in mESC showing that LH depletion can lead to both an increase and a decrease in gene expression (57), and that LH depletion in Drosophila can down regulate euchromatin genes (58).

As our nucleosome-resolution modeling study at the gene level reveals, the epigenetic regulation of chromatin by transcription factors is a complex dance that depends on the delicate composition of other internal and external factors present in the cell. Thus TFs collaborate with epigenetic marks and intrinsic chromatin features like linker lengths and nucleosome free regions to regulate the folding of chromatin at multiple scales and hence gene expression. Because the microdomains and loops affect access to and topologies of transcription start sites, these structural effects translate into effective regulation of cell differentiation and development, as well as human disease progression. Of course, genome regulation in the live cell involves much more than transcription factors. Other chromatin activators and repressors, architectural proteins, and RNAs are also involved. Although our implicit modeling of TF binding is a simple strategy, it provides clear trends and guiding insights into the regulation of chromatin architecture by protein binding. Further modeling with explicit proteins or consideration of various biomolecules will undoubtedly increase our understanding of genome regulation on the many structural and temporal levels involved in complex cellular contexts.

## 6 DATA AVAILABILITY

Matlab scripts for data analysis, as well as representative structures (in pdb format) for the 26 bp, 44 bp, 62 bp, and Life-Like fibers, and the Eed system are deposited on the Zenodo repository under https://zenodo.org/record/8199993.

## 7 AUTHOR CONTRIBUTIONS

S. P-L. designed research, performed simulations and analyses, wrote the original draft, and reviewed and edited the article. W. S. Chung performed simulations and analyses. J. H. performed simulations and analyses. T. S. designed research, reviewed and edited the article, acquired funding, and managed the project.

## ACKNOWLEDGMENTS

This work builds upon initial explorations of protein binding to chromatin fibers in our chromatin code by Dr. Gavin Bascom. Support from the National Institutes of Health, National Institute of General Medical Sciences Award R35-GM122562, National Science Foundation RAPID Award (2030377) from the Division of Mathematical Sciences, and Philip-Morris USA Inc to T.S. is gratefully acknowledged. The authors thank the HPC team at NYU for providing computational resources on the Greene NYU Supercomputer. S. P-L. is grateful for financial support from the Simons Center for Computational Physical Chemistry at NYU.

