## Supplementary material for "Regulation of Chromatin Architecture by Transcription Factor Binding": Figure SX, Table S1

### Supporting Information for "Regulation of Chromatin Architecture by Transcription Factor Binding"

**Table S1.** Setup of the Eed gene system. For each core is indicated the length of the linker DNA, if it has LH bound, and if its associated linker DNA has a Myc:Max binding site.

| Core # | Linker DNA | | LH | Myc:Max |
|---|---|---|---|---|
| | Beads | bp | | |
| 1 | 2 | 26 | 1 | 0 |
| 2 | 2 | 26 | 0 | 0 |
| 3 | 7 | 70 | 1 | 0 |
| 4 | 8 | 80 | 1 | 0 |
| 5 | 4 | 44 | 1 | 0 |
| 6 | 36 | 326 | 0 | 0 |
| 7 | 7 | 70 | 0 | 0 |
| 8 | 8 | 80 | 0 | 0 |
| 9 | 2 | 26 | 1 | 0 |
| 10 | 5 | 53 | 0 | 0 |
| 11 | 2 | 26 | 0 | 0 |
| 12 | 2 | 26 | 0 | 0 |
| 13 | 2 | 26 | 0 | 0 |
| 14 | 3 | 35 | 0 | 0 |
| 15 | 7 | 70 | 0 | 0 |
| 16 | 3 | 35 | 0 | 0 |
| 17 | 3 | 35 | 0 | 0 |
| 18 | 3 | 35 | 1 | 0 |
| 19 | 2 | 26 | 0 | 0 |
| 20 | 7 | 70 | 0 | 0 |
| 21 | 8 | 80 | 0 | 0 |
| 22 | 2 | 26 | 1 | 0 |
| 23 | 6 | 62 | 1 | 0 |
| 24 | 5 | 53 | 0 | 0 |
| 25 | 3 | 35 | 1 | 0 |
| 26 | 8 | 80 | 1 | 0 |
| 27 | 2 | 26 | 1 | 0 |
| 28 | 5 | 53 | 0 | 0 |
| 29 | 3 | 35 | 0 | 0 |
| 30 | 3 | 35 | 0 | 0 |
| 31 | 5 | 53 | 1 | 0 |

| | | | | |
|---|---|---|---|---|
| 32 | 7 | 70 | 1 | 0 |
| 33 | 2 | 26 | 0 | 0 |
| 34 | 5 | 53 | 0 | 0 |
| 35 | 4 | 44 | 0 | 0 |
| 36 | 4 | 44 | 0 | 0 |
| 37 | 2 | 26 | 0 | 0 |
| 38 | 3 | 35 | 0 | 0 |
| 39 | 7 | 70 | 1 | 0 |
| 40 | 2 | 26 | 1 | 0 |
| 41 | 5 | 53 | 0 | 0 |
| 42 | 3 | 35 | 1 | 0 |
| 43 | 7 | 70 | 1 | 0 |
| 44 | 2 | 26 | 0 | 0 |
| 45 | 2 | 26 | 0 | 0 |
| 46 | 2 | 26 | 0 | 0 |
| 47 | 5 | 53 | 0 | 0 |
| 48 | 5 | 53 | 1 | 0 |
| 49 | 2 | 26 | 1 | 0 |
| 50 | 4 | 44 | 1 | 0 |
| 51 | 3 | 35 | 1 | 0 |
| 52 | 2 | 26 | 1 | 0 |
| 53 | 2 | 26 | 0 | 0 |
| 54 | 6 | 62 | 0 | 0 |
| 55 | 2 | 26 | 0 | 0 |
| 56 | 36 | 326 | 0 | 0 |
| 57 | 2 | 26 | 0 | 0 |
| 58 | 2 | 26 | 0 | 0 |
| 59 | 5 | 53 | 1 | 0 |
| 60 | 2 | 26 | 1 | 0 |
| 61 | 6 | 62 | 0 | 0 |
| 62 | 3 | 35 | 0 | 0 |
| 63 | 8 | 80 | 0 | 0 |
| 64 | 6 | 62 | 1 | 0 |
| 65 | 6 | 62 | 0 | 0 |
| 66 | 6 | 62 | 0 | 0 |
| 67 | 8 | 80 | 0 | 0 |
| 68 | 5 | 53 | 0 | 0 |
| 69 | 4 | 44 | 0 | 0 |
| 70 | 2 | 26 | 0 | 0 |
| 71 | 3 | 35 | 0 | 0 |
| 72 | 3 | 35 | 0 | 0 |
| 73 | 4 | 44 | 1 | 0 |
| 74 | 5 | 53 | 0 | 0 |
| 75 | 2 | 26 | 0 | 0 |
| 76 | 8 | 80 | 0 | 0 |
| 77 | 2 | 26 | 0 | 0 |
| 78 | 18 | 168 | 1 | 0 |
| 79 | 3 | 35 | 1 | 0 |
| 80 | 6 | 62 | 0 | 0 |
| 81 | 5 | 53 | 0 | 0 |

| | | | | |
|---|---|---|---|---|
| 82 | 4 | 44 | 0 | 0 |
| 83 | 2 | 26 | 0 | 0 |
| 84 | 5 | 53 | 0 | 0 |
| 85 | 3 | 35 | 0 | 0 |
| 86 | 2 | 26 | 1 | 0 |
| 87 | 2 | 26 | 1 | 0 |
| 88 | 8 | 80 | 0 | 0 |
| 89 | 7 | 70 | 1 | 0 |
| 90 | 6 | 62 | 1 | 0 |
| 91 | 8 | 80 | 1 | 0 |
| 92 | 18 | 168 | 1 | 0 |
| 93 | 2 | 26 | 1 | 0 |
| 94 | 5 | 53 | 0 | 0 |
| 95 | 4 | 44 | 0 | 0 |
| 96 | 3 | 35 | 1 | 0 |
| 97 | 18 | 168 | 0 | 0 |
| 98 | 3 | 35 | 0 | 0 |
| 99 | 8 | 80 | 0 | 0 |
| 100 | 4 | 44 | 1 | 1 |
| 101 | 18 | 168 | 0 | 1 |
| 102 | 4 | 44 | 0 | 1 |
| 103 | 4 | 44 | 0 | 0 |
| 104 | 3 | 35 | 1 | 0 |
| 105 | 2 | 26 | 1 | 0 |
| 106 | 3 | 35 | 0 | 0 |
| 107 | 6 | 62 | 0 | 0 |
| 108 | 6 | 62 | 1 | 0 |
| 109 | 3 | 35 | 1 | 0 |
| 110 | 18 | 168 | 1 | 0 |
| 111 | 4 | 44 | 1 | 0 |
| 112 | 2 | 26 | 1 | 0 |
| 113 | 4 | 44 | 0 | 0 |
| 114 | 4 | 44 | 1 | 0 |
| 115 | 6 | 62 | 0 | 0 |
| 116 | 4 | 44 | 0 | 0 |
| 117 | 5 | 53 | 0 | 0 |
| 118 | 7 | 70 | 1 | 0 |
| 119 | 2 | 26 | 0 | 0 |
| 120 | 5 | 53 | 0 | 0 |
| 121 | 8 | 80 | 1 | 0 |
| 122 | 2 | 26 | 1 | 0 |
| 123 | 2 | 26 | 0 | 1 |
| 124 | 2 | 26 | 0 | 1 |
| 125 | 4 | 44 | 0 | 1 |
| 126 | 2 | 26 | 0 | 1 |
| 127 | 3 | 35 | 0 | 1 |
| 128 | 2 | 26 | 0 | 1 |
| 129 | 5 | 53 | 0 | 1 |

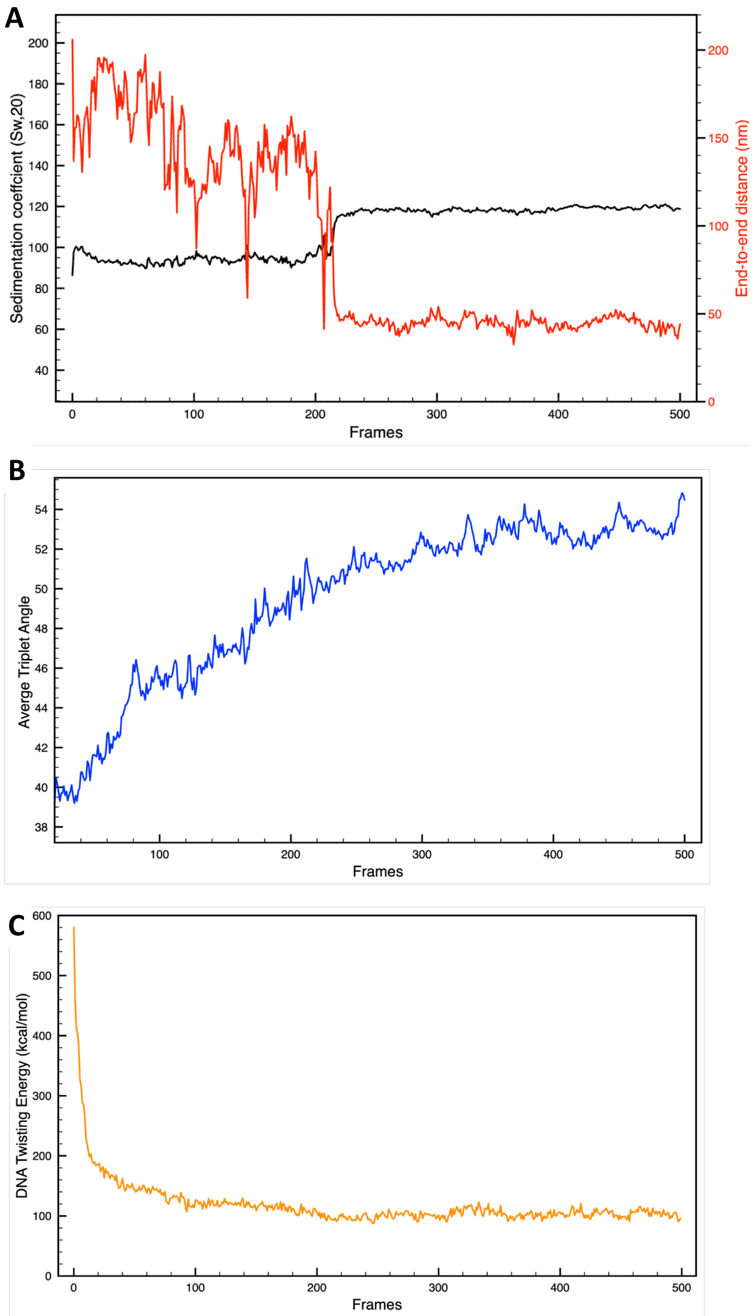


**Figure S1. Convergence check. A)** Sedimentation coefficient and end-to-end distance evolution; **B)** average triplet angle evolution, and **C)** sample energy evolution for the 26 bp system with 5 TF binding regions.

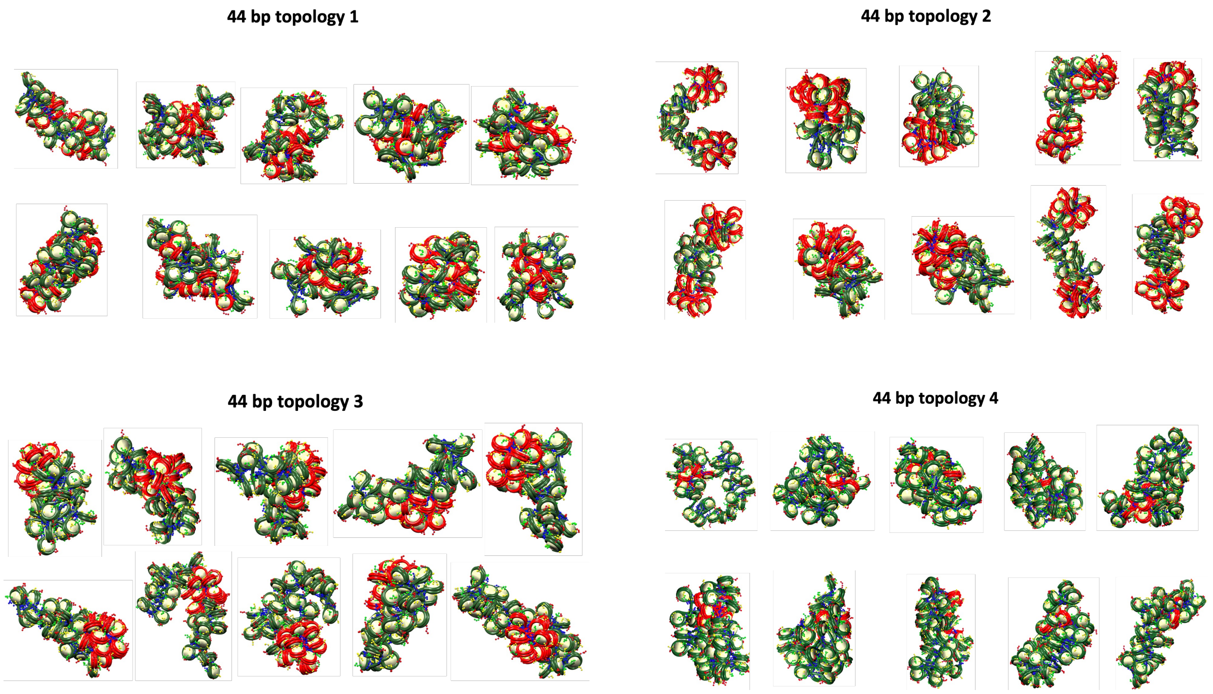


**Figure S2.** Final configurations for each of the 10 trajectories of the 44 bp system with four different TF binding topologies.

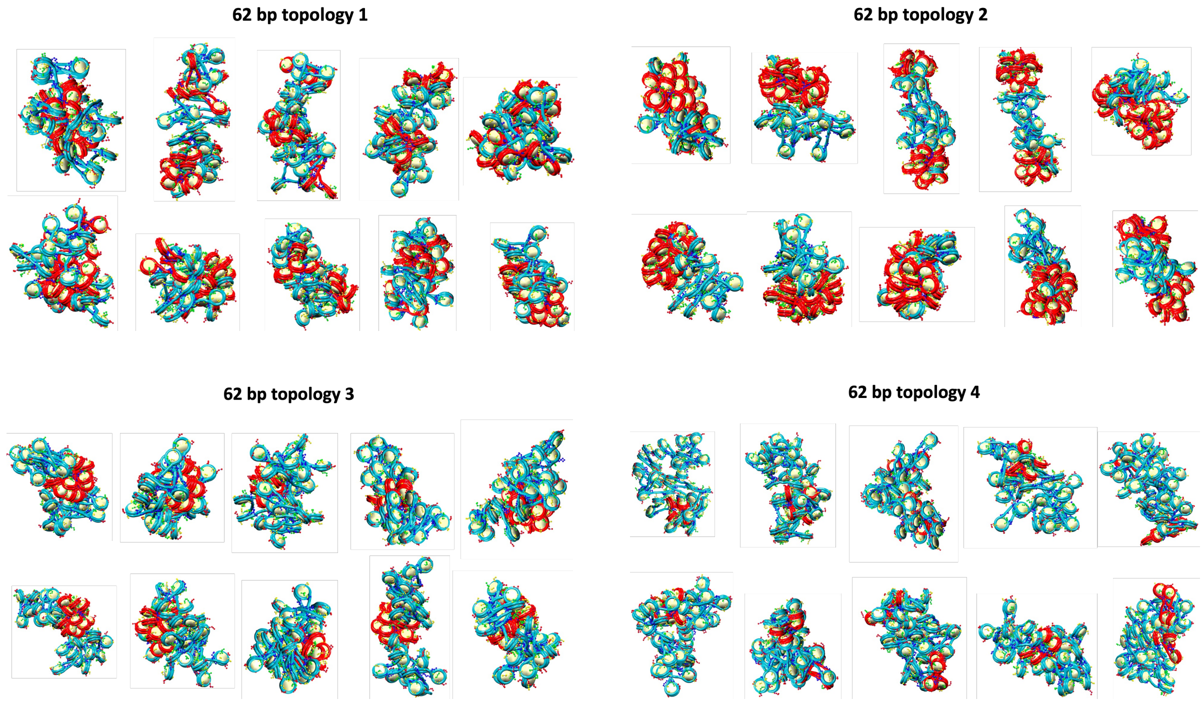


**Figure S3.** Final configurations for each of the 10 trajectories of the 62 bp system with four different TF binding topologies.

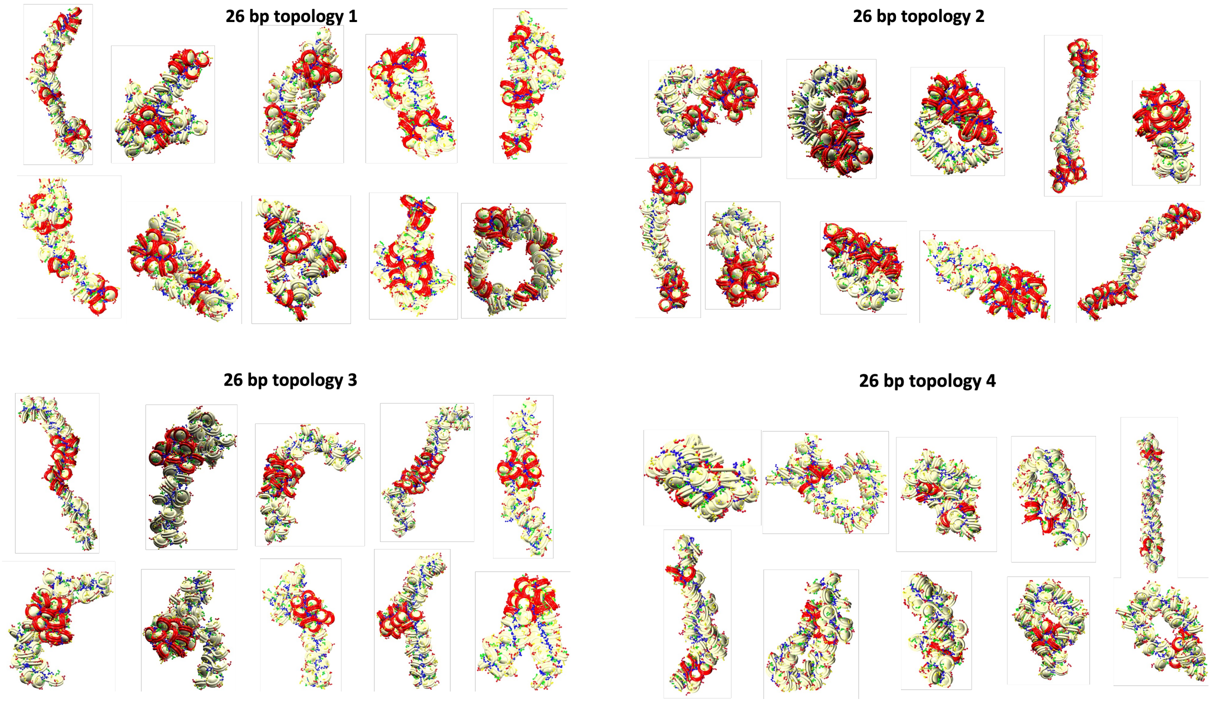


**Figure S4.** Final configurations for each of the 10 trajectories of the 26 bp system with four different TF binding topologies.

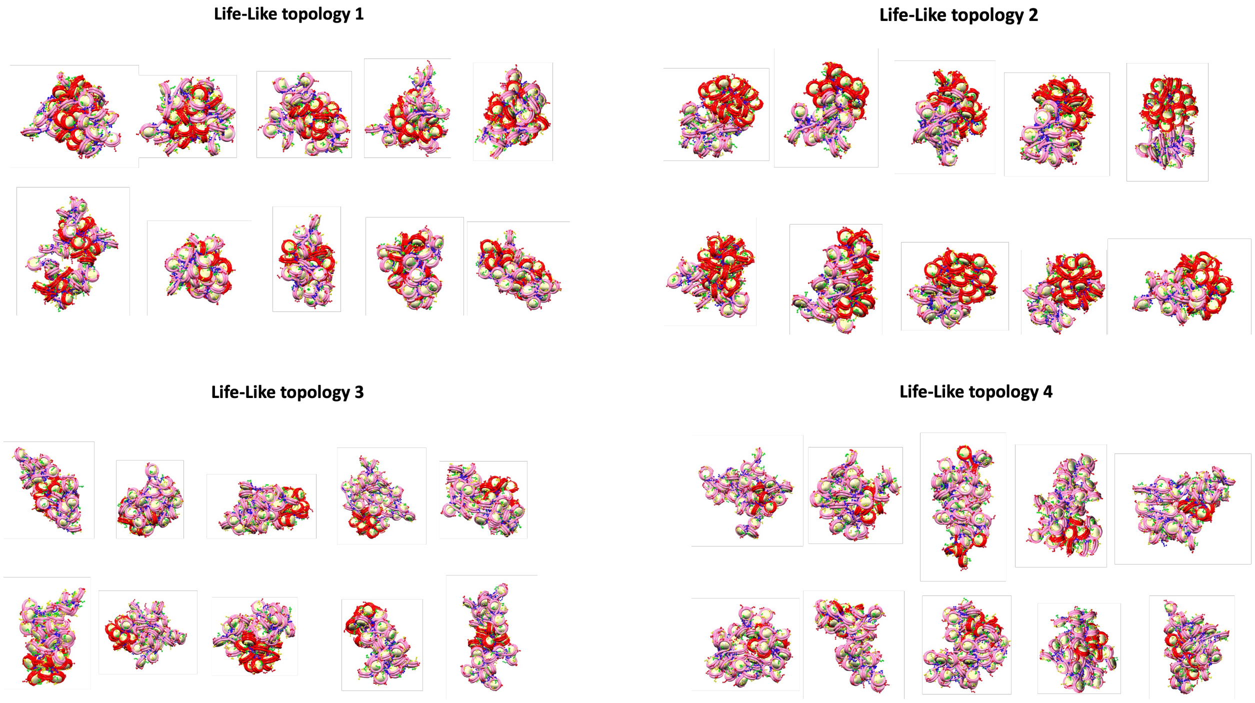


**Figure S5.** Final configurations for each of the 10 trajectories of the Life-Like system with four different TF binding topologies.

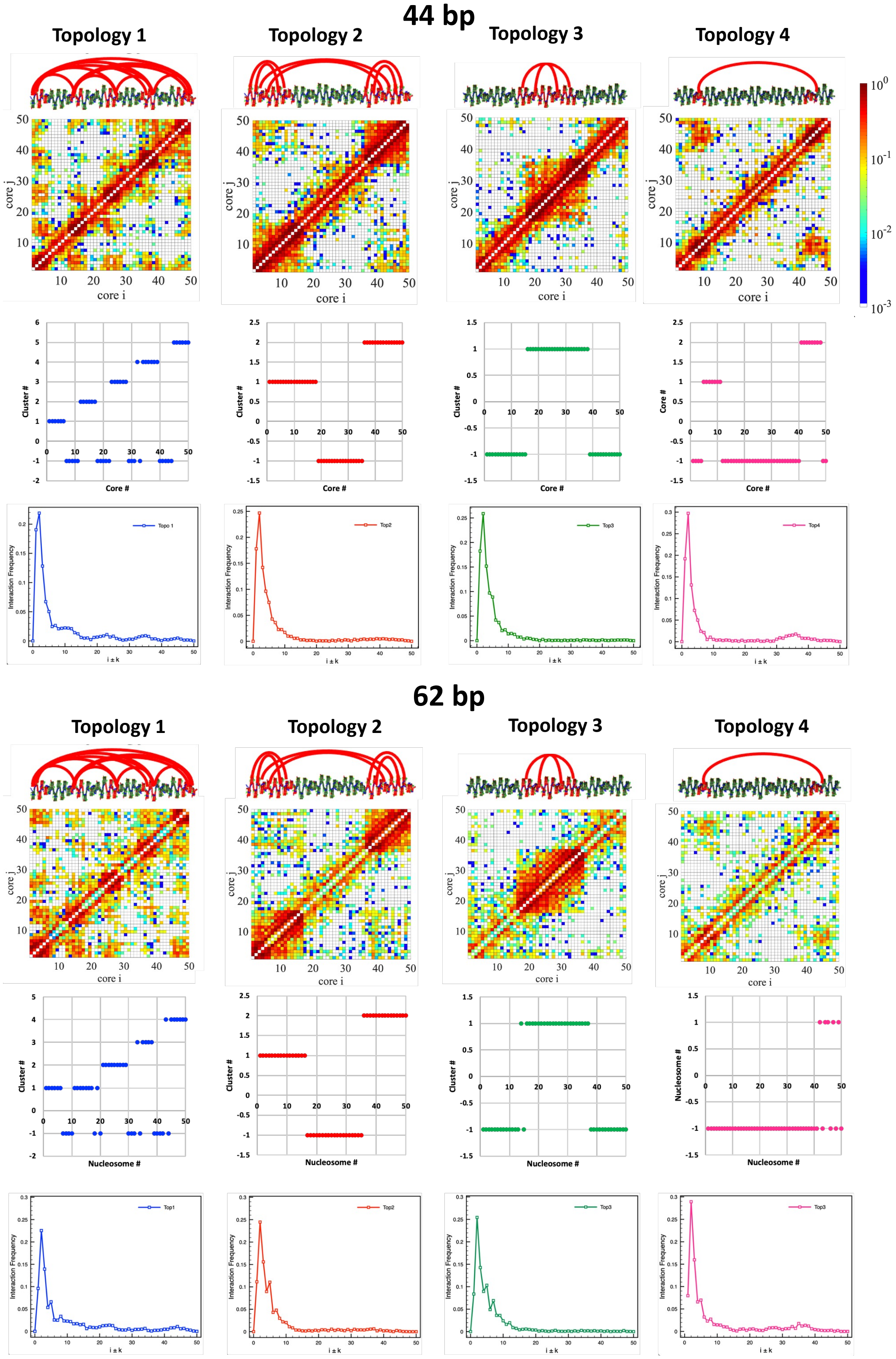

44 bp
Topology 1
Topology 2
Topology 3
Topology 4
core j
core i
Cluster #
Core #
Interaction Frequency
i ± k
Topo 1
Top2
Top3
Top4
62 bp
Topology 1
Topology 2
Topology 3
Topology 4
Nucleosome #
Top1
Top2
Top3
Top3


**Figure S6.**

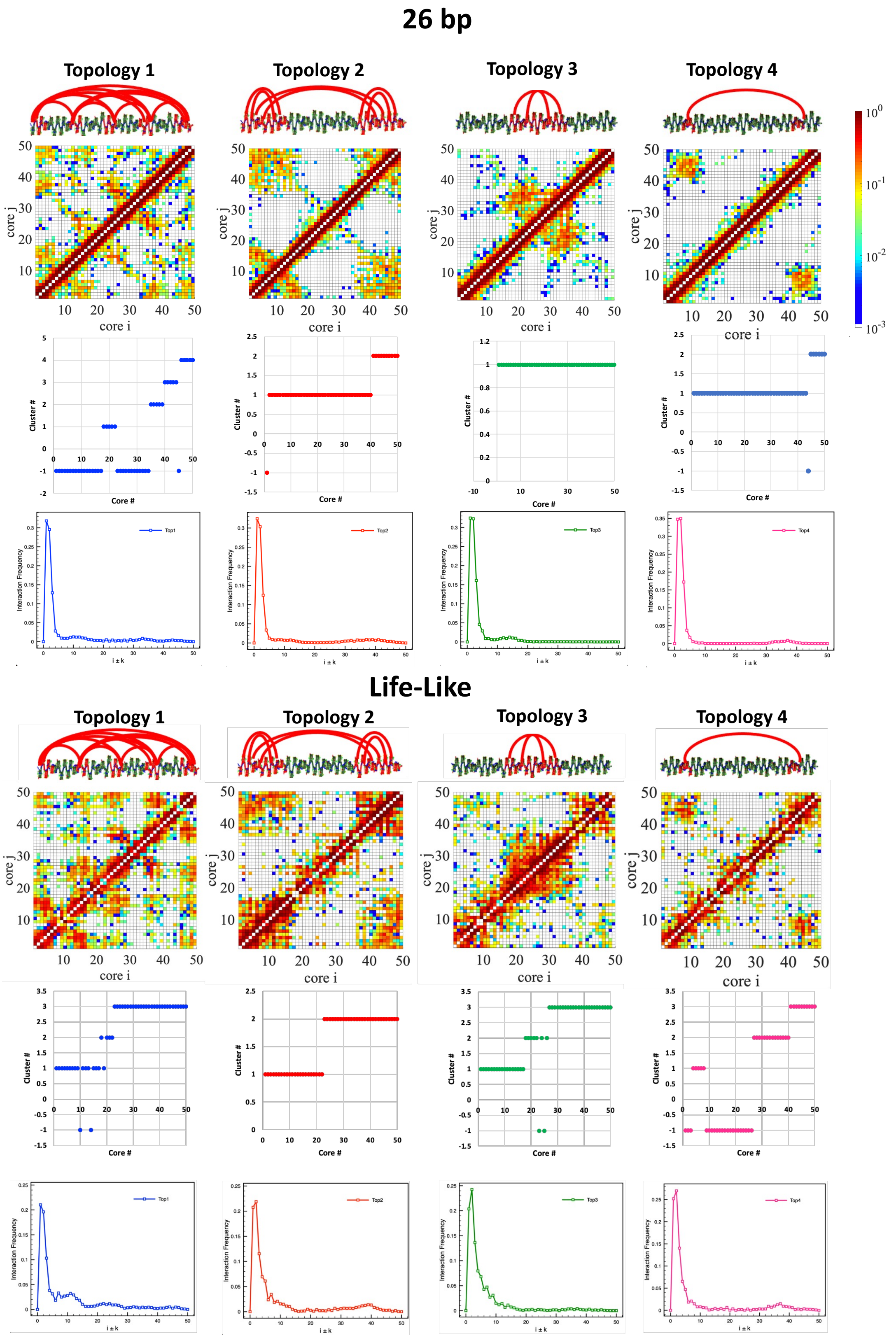


**Figure S6 cont.**

**Figure S6.** DBSCAN clustering supports the formation of microdomains. DBSCAN clustering (1) was performed on distance matrices obtained from the contact maps with nucleosome resolution ($d = 1/freq$) and using minpoint = 5 and epsilon = 3, 2, 2, and 1.4 for the 62 bp, 44 bp, Life-Like, and 26 bp systems, respectively. Internucleosome interactions in one dimension are plotted as $Int. Freq. vs. i \pm k$.

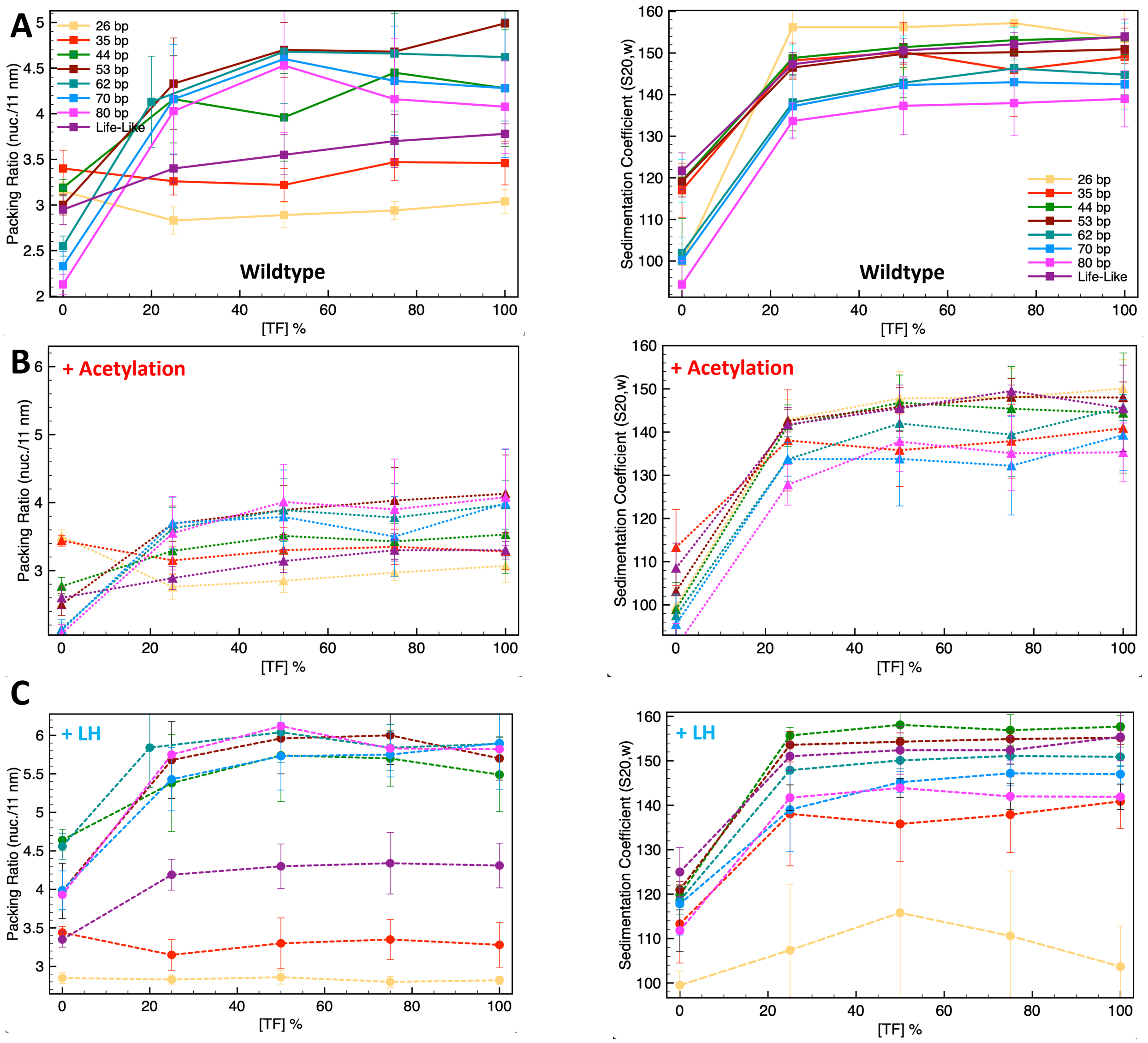


**Figure S7.** TF saturation curves for the uniform systems of 26, 35, 44, 53, 62, 70, and 80 bp, as well as for the life-like fiber non uniform system.

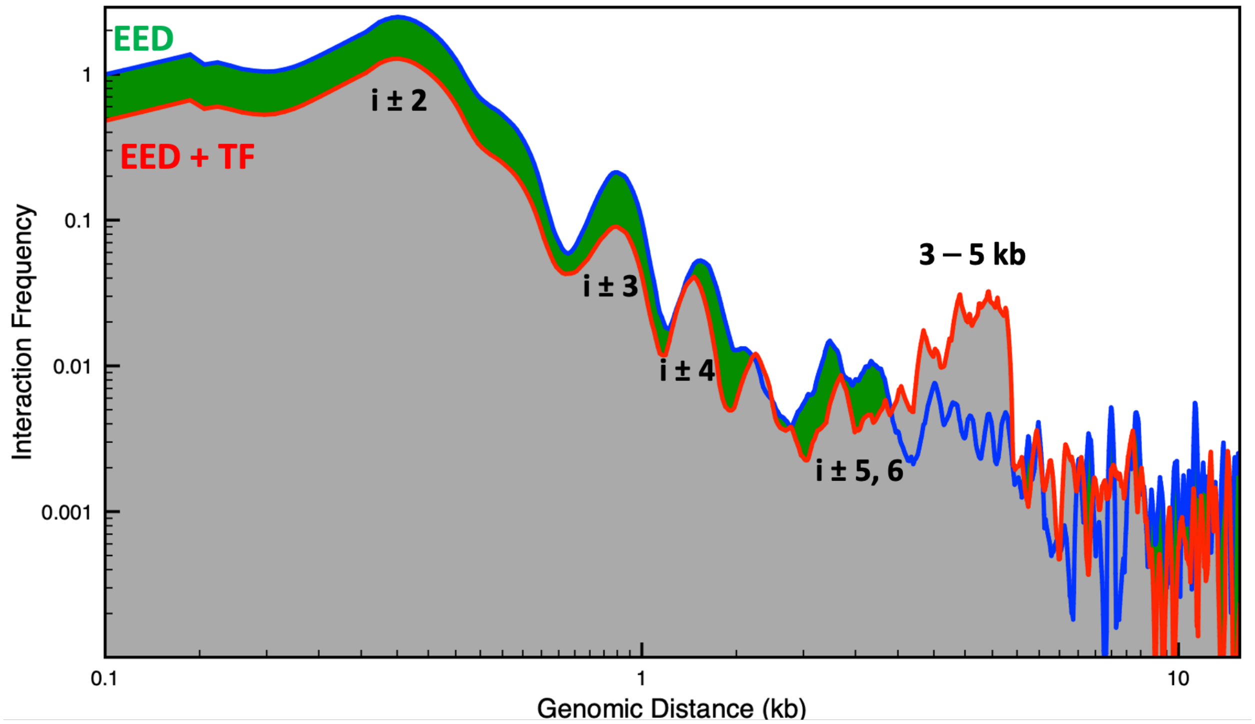


**Figure S8.** Genomic interaction frequency versus genomic distance for the EED gene with and without TF bound.

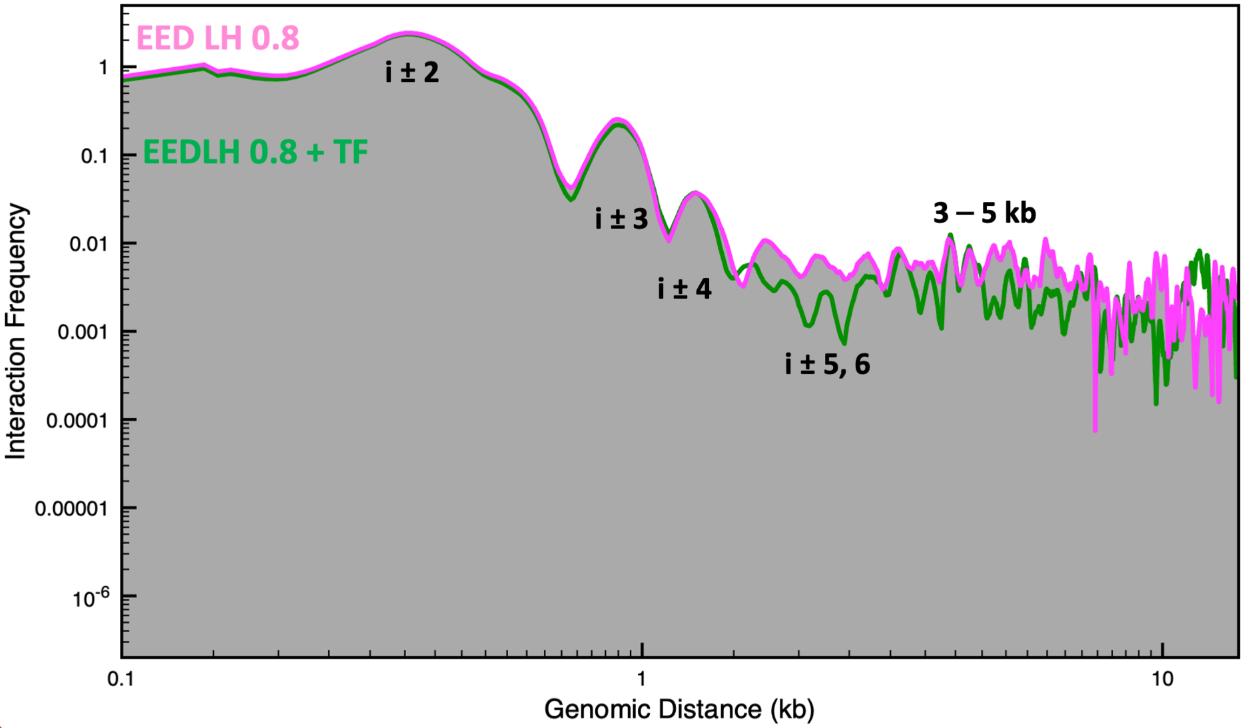


**Figure S9.** Genomic interaction frequency versus genomic distance for the EED gene with an LH density of 0.8 with and without TF bound.
